# Dimensions of anxiety and depression and neurophysiological indicators of error-monitoring: Relationship with delta and theta oscillatory power and error-related negativity amplitude

**DOI:** 10.1101/872119

**Authors:** Alexandra M. Muir, Ariana Hedges-Muncy, Ann Clawson, Kaylie A. Carbine, Michael J. Larson

## Abstract

Error-monitoring processes may be affected by transdiagnostic dimensions of psychopathology symptoms including trait anxiety, worry, and severity of depressive symptoms. We tested the relationship between continuous measures of anxiety and depressive symptomology and neural correlates of error-monitoring as measured by time-frequency domain delta and theta oscillatory power and time domain error-related negativity (ERN) amplitude extracted from the electroencephalogram (EEG). Secondary analyses tested for diagnostic group differences in error-related neural responses in individuals with generalized anxiety disorder (GAD), major depressive disorder (MDD), and comorbid psychiatric disorders. 178 participants (104 female, *M[SD]_age_* = 21.7[4.6]) with a wide range of psychopathology symptoms completed a modified version of the Eriksen flanker task and symptom questionnaires. Residualized difference values between correct and error trials for delta/theta power and error/correct ERN amplitude were dependent variables. Linear regression analyses adjusted for age and sex showed nonsignificant associations of symptom dimension measures with error-related residualized delta/theta power or residualized ERN amplitude. Subset analyses on those with confirmed psychopathology diagnoses also did not predict residualized error-related delta/theta power nor ERN amplitude. Exploratory analyses with only error trial delta/theta power and ERN amplitude also revealed nonsignificant relationships. Taken in the context of previous literature, results suggest a heterogeneous relationship between depressive and anxiety symptom dimensions and neurophysiological indices of error-monitoring.

**Impact Statement:** In line with the RDoC framework, we tested the relationship between anxiety and depressive symptom dimensions and neural indices of error-processing (delta and theta power, error-related negativity ERP amplitude) in 178 participants with a range of pathology symptoms. A non-significant relationship emerged between neural and symptom measures suggesting anxiety and depressive symptomology have a nuanced relationship with error-monitoring in a large sample across a range of anxiety and depression symptoms.

## 1 Introduction

Error-monitoring, an individual’s ability to detect an incorrect response and subsequently adjust to improve future behavior, is an essential skill to achieve successful goal-directed behavior (Mohamed, Börger, Geuze, & van der Meere, 2019; Rabbitt & Rodgers, 1977). Individual differences in error-monitoring may be related to personality traits and transdiagnostic psychopathology symptoms, such as worry, negative affect, impulsivity, and conscientiousness (Hill, Samuel, & Foti, 2016; Moser, Moran, & Jendrusina, 2012). Error-monitoring processes are altered in individuals with psychopathology. For example, individuals with generalized anxiety disorder and obsessive-compulsive disorder have heightened error-monitoring processes (Riesel, Kathmann, & Endrass, 2014; Weinberg, Olvet, & Hajcak, 2010); however, the relationship between error-monitoring processes and symptoms of psychopathology is heterogeneous for other disorders such as major depressive disorder (Aarts, Vanderhasselt, Otte, Baeken, & Pourtois, 2013; Gorka & Phan, 2017; Weinberg, Liu, & Shankman, 2016). We tested the relationship between transdiagnostic symptom dimensions of depression and anxiety and neurophysiological reflections of error-monitoring processes, including event-related potentials (ERP) and electroencephalogram (EEG) oscillatory power in a sample with a wide range of psychopathology symptoms. A secondary aim was to test diagnostic group differences in neurophysiological responses to errors in individuals with confirmed diagnoses of generalized anxiety disorder (GAD), major depressive disorder (MDD), and comorbid psychiatric disorders.

### 1.1 Neurophysiological Measures of Error Monitoring

The error-related negativity (ERN) is an ERP often used to quantify neural manifestations of error-monitoring. The ERN is a negative deflection in the ERP waveform approximately 0 to 150 ms following an erroneous response (Gehring, Goss, Coles, Meyer, & Donchin, 1993) that originates in the anterior cingulate cortex (ACC; van Veen & Carter, 2002). Despite numerous theories concerning the functional significance of the ERN, the current consensus is that the ERN represents an early monitoring system interpreting cognitive or emotional responses to errors, after which additional cognitive resources are recruited to improve future behavior (Gehring et al., 1993; Larson, Clayson, & Clawson, 2014; Proudfit, Inzlicht, & Mennin, 2013; Weinberg, Liu, Hajcak, & Shankman, 2015).

In addition to time domain measures, analyses of EEG data in the time and frequency domains can be used to quantify neural response to errors. Time-frequency analyses measure the magnitude of frequency band oscillations and are thought to reflect increased synchronization of a group of neurons working together to produce a cognitive response (Buzaski, 2006). While time domain measures such as ERN amplitude capture phase-locked data, time-frequency measures capture both phase- and non-phase locked data, resulting in a richer representation of the EEG signal (Cohen, 2014). Thus, utilization of both time and time-frequency measures to quantify neural response to errors provides a rich and holistic view of the neurophysiological processes related to error-monitoring.

Oscillations in the delta (1-3 Hz) and theta (4-8 Hz) frequency bands are thought to reflect error-monitoring processes. Specifically, both midline delta and theta activity increase directly following an incorrect response compared to following correct responses (Cavanagh, Cohen, & Allen, 2009; Luu & Tucker, 2001; Munneke, Nap, Schippers, & Cohen, 2015) and are present in frequency decompositions of the ERN (Luu & Tucker, 2001; Yordanova, Falkenstein, Hohnsbein, & Kolev, 2004). The functional roles of delta- and theta-band activity may also be dissociable (Cohen & Cavanagh, 2011), with evidence suggesting that the delta-band is primarily associated with error-monitoring, while theta-band activity includes both conflict-(i.e., the simultaneous presentation of competing options) and error-related processes (Cohen & Cavanagh, 2011). The ERN and delta/theta oscillatory power quantify both similar and independent portions of neural signal (Cavanagh, Meyer, & Hajcak, 2017; Munneke et al., 2015), suggesting the utility of using both the ERP and oscillation-based measures to quantify neural indices of error-monitoring.

### 1.2 Symptoms of Anxiety and Depression and Error-Monitoring Processes

There is increasing focus on the relationship between symptom dimensions of psychopathology on a continuous scale and error-monitoring processes, regardless of formal psychiatric diagnosis (i.e., a transdiagnostic approach). This approach is in line with the Research Domain Criteria (RDoC) initiative, which aims to establish cognitive and behavioral constructs under which psychopathology can be studied, regardless of traditional diagnostic labels. In the past, diagnostic status was used to group individuals, after which those group differences in error-monitoring were tested (i.e., Aarts et al., 2013; Weinberg et al., 2010). However, it is possible traditional nosology of psychopathology may not be valid nor capture underlying aberrant biology that results in presentation of abnormal behavior (Cuthbert & Insel, 2013). Error-monitoring processes fit well in the RDoC framework, as error-monitoring has the potential to link psychopathology to underlying deviant neural functioning that affects outward behavior (Hanna & Gehring, 2016). Thus, investigating relationships between transdiagnostic symptom dimensions and personality traits in samples with a wide range of psychopathology symptoms allows for a better understanding of what factors may influence individual differences in error-monitoring abilities. The primary approach employed in the current study was to use individual difference psychopathology symptom measures to test for relationships with error-related neurophysiology, regardless of psychiatric diagnosis.

Trait anxiety is a stable personality trait in which an individual tends to be in a continuously anxious state (Kennedy, Schwab, Morris, & Beldia, 2001). Individuals high on trait anxiety show greater frontal midline theta when compared to those lower on scales of trait anxiety (Schmidt, Kanis, Holroyd, Miltner, & Hewig, 2018); however, other results suggest no relationship between trait anxiety and midline theta (Neo & McNaughton, 2011). In the time domain, larger ERN component amplitude is related to higher trait anxiety scores (Olvet & Hajcak, 2008), which has been interpreted as an indicator of greater expectancy violation in individuals with higher trait anxiety (Compton et al., 2007). When examining the relationship between anxiety symptoms and neural mechanisms of error-monitoring, it is important to dissociate state anxiety from trait anxiety. Anxiety inducing paradigms produced no change in ERN amplitude (Moser, Hajcak, & Simons, 2005) suggesting the ERN is trait-like in nature (Olvet & Hajcak, 2008). Therefore, in the current study, only the trait subscale of the State Trait Anxiety Inventory (STAI) was used to investigate the relationship between trait anxiety and neural measures of error-monitoring.

Along with trait anxiety, anxious apprehension (i.e., worry), depressive symptomology, and biological sex may be factors influencing the neurophysiological representations of error-monitoring processes. Anxious apprehension (i.e., worry) is a cognitive component of anxiety where worrisome thoughts dominate day to day life (Nitschke, Heller, Imig, McDonald, & Miller, 2001). Individuals who scored high on measures of anxious apprehension, regardless of diagnosis, displayed enhanced ERN amplitude when compared to controls (Hajcak, McDonald, & Simons, 2003; Moser et al., 2012; Moser, Moran, Kneip, Schroder, & Larson, 2016; Moser, Moran, Schroder, Donnellan, & Yeung, 2013). When looking at the relationship between depressive symptomology and error monitoring processes, there is great heterogeneity in the literature with some evidence that ERN amplitude is not related to depressive symptoms (Chang, Davies, & Gavin, 2010; Schroder, Moran, Infantolino, & Moser, 2013). Other research indicates that ERN amplitude is related to facets of melancholia (Weinberg et al., 2016), suggesting that the ERN may be more specifically related to facets of depression rather than depressive symptoms as a whole. In addition to the possible modulation of ERN amplitude by worry and depressive symptomology, ERN amplitude may differ as a function of biological sex; however the current literature is unclear as to whether men or women display greater ERN amplitudes (Fischer, Danielmeier, Villringer, Klein, & Ullsperger, 2016; Hill, Ait Oumeziane, Novak, Rollock, & Foti, 2018; Larson, South, & Clayson, 2011; Moser et al., 2016). Thus, it is important to account for biological sex when examining individual differences in error-monitoring.

### 1.3 Diagnostic Status and Error-Monitoring Processes

There is a significant amount of heterogeneity in studies of error-monitoring processes within individuals formally diagnosed with MDD, GAD, or comorbid disorders. ERN amplitude is generally heightened in individuals with anxiety disorders such as GAD (Meyer, Nelson, Perlman, Klein, & Kotov, 2018; Weinberg, Liu, et al., 2015) and enhanced theta power reliably dissociated individuals with GAD from psychiatrically healthy controls (Cavanagh et al., 2017). However, there is also evidence that ERN amplitude is unchanged in people with diagnosed GAD (Kujawa et al., 2016; Xiao et al., 2011). In individuals with comorbid anxiety and depression, there is evidence that ERN amplitude is unchanged from psychiatrically healthy controls (Weinberg, Klein, & Hajcak, 2012; Weinberg, Kotov, & Proudfit, 2015), implying comorbid depression may moderate the relationship between ERN amplitude and GAD diagnostic status. When looking at individuals diagnosed with MDD, although there is some evidence that individuals with MDD have an enhanced ERN when compared to controls (Aarts et al., 2013; Chiu & Deldin, 2007; Holmes & Pizzagalli, 2008, 2010), other evidence suggests that either ERN amplitude is blunted (Olvet, Klein, & Hajcak, 2010; Weinberg et al., 2016), or that there is no difference in ERN amplitude between those with MDD and those without (Gorka & Phan, 2017; Moran, Schroder, Kneip, & Moser, 2017; Weinberg et al., 2012). In addition to this heterogeneity of evidence, there is a lack of evidence present concerning error-related delta and theta power in relation to GAD, MDD, and comorbid disorders. As such, combining time domain and time-frequency domain measures of error processing in GAD, MDD, and comorbid disorders may assist in elucidating the relationship between diagnostic status and neural indices of error-monitoring.

### 1.4 Aims and Hypotheses

The current study had two aims. Our primary aim was to quantify the relationship between symptom dimensions of trait anxiety, worry, and depressive symptomology and error-monitoring processes in individuals with a wide range of symptoms regardless of psychiatric diagnosis using commonly utilized measures of psychopathology. To isolate error-related activity instead of general response-related activity, residualized difference values between correct and error trials for delta/theta power and ERN amplitude were used as the dependent variable of interest (Meyer, Lerner, Reyes, Laird, & Hajcak, 2017). We hypothesized, based on the current literature, that higher trait anxiety and worry would be related to residual delta power, theta power, and ERN amplitude. Due to the heterogeneity of the literature, we also hypothesized there would be no relationship between depressive symptoms and residual delta power, theta power, and ERN amplitude. A secondary aim of the current study was to characterize error-monitoring processes in individuals with a diagnosis of GAD, MDD, or comorbid disorders. Similar to the previous hypotheses, we hypothesized that individuals with GAD would have greater residualized delta power, theta power, and ERN amplitude when compared to psychiatrically healthy controls, but there would be no difference in dependent variables between those with MDD, comorbid disorders, and controls.

## 2 Method

All data and code are posted on the Open Science Framework (OSF) and can be found at https://osf.io/pujsv. All methods are in compliance with the methodological reporting checklist for EEG/ERP data as outlined in Keil et al. (2014; see also Clayson, Carbine, Baldwin, & Larson, 2019). A subset of the current data testing different data aspects and hypotheses have been previously published (see Baldwin, Larson, & Clayson, 2015; Clawson, Clayson, & Larson, 2013).

### 2.1 Participants

Procedures were approved by the Brigham Young University Institutional Review Board. Psychiatrically-healthy control participants were recruited through undergraduate psychology courses, whereas individuals with psychiatric diagnoses and elevated symptoms of psychopathology were recruited through flyers placed at the local university counseling center and community mental health centers. All participants were compensated through course credit or monetary payment.

The final sample consisted of 178 participants (female = 104; *M(SD)_age_* = 21.7[4.6]). For those with psychopathology, diagnoses were initially made by a psychiatrist, psychologist, or physician in the community and subsequently confirmed upon enrollment using the Mini-International Neuropsychiatric Inventory (MINI; Sheehan et al., 1998). The MINI has a high concordance rate with the Structured Clinical Interview for DSM-IV Axis I disorders (SCID) but requires less time to administer (Sheehan et al., 1998). Participants were excluded if they had medication changes within the two months prior to data collection, they had a diagnosis of a psychotic or bipolar disorder, they reported a learning disorder or attention deficit/hyperactivity disorder, they had a history of substance use or dependence, neurological disease, or they were left-handed. At the time of participation 33.1% of all participants were taking a psychotropic medication (GAD = 78.6%, MDD = 60.7%, Comorbid = 58.8% [see Table S1 in the supplementary material on OSF for a list of comorbid disorders] No confirmed diagnosis = 61.8%, Control = 0%). The proportion of participants taking psychotropic medications did not differ between the three (GAD, MDD, comorbid) psychopathology groups (*χ^2^*(2) = 1.62, *p* = 0.44; see Table S2 in the supplementary material on OSF).

**Table 1.** Means and standard deviations for demographics and questionnaires

|  | Group | Mean | Standard<br>Deviation | Range<br>(min,max) | Chronbach's<br>Alpha |
| --- | --- | --- | --- | --- | --- |
| Age | Overall | 21.67 | 4.59 | 18,53 |  |
|  | GAD | 22.07 | 4.53 | 18, 32 |  |
|  | MDD | 21.89 | 2.15 | 18,26 |  |
|  | Comorbid | 22.53 | 6.56 | 18,47 |  |
|  | Control | 20.61 | 2.27 | 18, 27 |  |
| BDI | Overall | 13.97 | 12.45 | 0,53 | 0.95 |
|  | GAD | 17.36 | 9.29 | 3,34 | 0.88 |
|  | MDD | 21.07 | 12.17 | 1,39 | 0.93 |
|  | Comorbid | 25.68 | 17.82 | 0,53 | 0.95 |
|  | Control | 5.85 | 4.57 | 0,21 | 0.81 |
| PSWQ | Overall | 40.68 | 12.68 | 4,67 | 0.93 |
|  | GAD | 46.57 | 8.4 | 32, 61 | 0.9 |
|  | MDD | 43.54 | 9.97 | 25, 58 | 0.92 |
|  | Comorbid | 50.29 | 8.51 | 34, 64 | 0.89 |
|  | Control | 34.99 | 13.66 | 4, 67 | 0.92 |
| STAI | Overall | 44.23 | 13.6 | 20, 74 | 0.95 |
|  | GAD | 52.93 | 10.83 | 29, 66 | 0.91 |
|  | MDD | 52.82 | 11.53 | 24, 68 | 0.91 |
|  | Comorbid | 59.35 | 8.65 | 45, 74 | 0.83 |
|  | Control | 34.42 | 7.29 | 20, 55 | 0.85 |

**Table 2.** Means and standard deviations for dependent variables and trial numbers.

|  | Group | Mean | Standard<br>Deviation | Range<br>(min,max) |
| --- | --- | --- | --- | --- |
| Error delta | Overall | 2.46 | 1.41 | -4.85, 6.79 |
|  | GAD | 2.07 | 1.46 | 0.22, 4.80 |
|  | MDD | 2.17 | 2.04 | -4.85, 6.45 |
|  | Comorbid | 2.39 | 1.41 | 0.48, 5.40 |
|  | Control | 2.65 | 1.62 | -1.35, 6.79 |
| Correct delta | Overall | 0.99 | 1.31 | 2.50, 8.59 |
|  | GAD | 0.94 | 1.08 | -1.21, 2.51 |
|  | MDD | 0.87 | 0.97 | -1.81, 2.34 |
|  | Comorbid | 0.98 | 0.86 | -0.40, 3.18 |
|  | Control | 0.89 | 1.33 | -2.50, 8.59 |
| Error theta | Overall | 1.95 | 2.06 | -8.67, 7.70 |
|  | GAD | 2.06 | 1.62 | -1.35, 4.40 |
|  | MDD | 1.80 | 2.70 | -8.67, 6.97 |
|  | Comorbid | 2.34 | 2.01 | 0.03, 5.43 |
|  | Control | 1.94 | 2.03 | -3.68, 7.70 |
| Correct Theta | Overall | -0.14 | 1.56 | -7.29, 11.87 |
|  | GAD | 0.17 | 1.09 | -2.15, 2.16 |
|  | MDD | -0.10 | 0.89 | -2.30, 1.15 |
|  | Comorbid | 0.18 | 1.02 | -1.44, 3.07 |
|  | Control | -0.46 | 1.42 | -7.29, 5.30 |
| Error-related negativity<br>(ERN) | Overall | -0.12 | 2.59 | -11.19, 7.29 |
|  | GAD | -0.18 | 1.56 | -2.28, 2.52 |
|  | MDD | -0.17 | 2.75 | -11.19, 4.11 |
|  | Comorbid | -0.42 | 1.78 | -3.06, 3.70 |
|  | Control | -0.03 | 2.81 | -9.51, 7.29 |
| Correct-related<br>negativity<br>(CRN) | Overall | 1.46 | 2.75 | -11.87, 18.38 |
|  | GAD | 1.06 | 1.10 | -0.76, 3.37 |
|  | MDD | 1.38 | 2.12 | -3.62, 5.02 |
|  | Comorbid | 1.08 | 1.66 | -1.42, 3.77 |
|  | Control | 1.78 | 3.17 | -11.87, 18.38 |
| Correct Trials | Overall | 648.7 | 114.62 | 164, 784 |
|  | GAD | 662.79 | 103.12 | 443, 771 |
|  | MDD | 629.82 | 140.68 | 192, 775 |
|  | Comorbid | 606.16 | 151.65 | 164,779 |
|  | Control | 652.81 | 102.16 | 278, 773 |
| Error Trials | Overall | 47.4 | 31.07 | 10, 170 |
|  | GAD | 49.36 | 29.52 | 14, 108 |
|  | MDD | 58.89 | 31.23 | 18, 136 |
|  | Comorbid | 49.47 | 32.15 | 14, 119 |
|  | Control | 40.13 | 28.69 | 11, 158 |

*Note:* Correct trials were randomly selected to match the number of error trials in the time-frequency data analyses
Delta/Theta power = db Change from Baseline
ERN/CRN = microvolts

Participants were excluded if they had ERP noise levels greater than 20 (root mean square of the residual noise after the consistent ERP is canceled by inverting every other trial; see Schimmel, 1967), if they had less than 50% accuracy on the computerized tasks, or if they had missing or incomplete questionnaire data. To ensure similar number of trials for the oscillatory power and ERP analyses, all participants had a minimum of ten useable trials for all conditions. Because reliability is a product of the context of a current sample and study (Clayson & Miller, 2017a) dependability of ERN amplitude (for both error and correct trials) was estimated using the ERP Reliability Analysis Toolkit in Matlab (Clayson & Miller, 2017b). This toolkit uses generalizability theory to estimate the g-theory reliability analogue known as dependability in ERP components. The error trials had an average dependability of 0.63 and the correct trials had an average dependability of 0.86.

Of the 178 participants, 32 participants who originally indicated they had a psychiatric diagnosis were excluded from secondary diagnostic analyses due to a lack MINI confirmation of their diagnostic status (i.e., they were diagnosed by a practitioner in the community, but the diagnosis was not confirmed by MINI administration in the lab). Thus, the final sample for our secondary diagnostic subgroup secondary analyses consisted of 146 participants (87 controls, 61 individuals with psychopathology; *M(SD)_age_* = 21.3(3.4), female = 87; n_GAD_ = 14; n_MDD_ = 28; n_Comorbid_ = 19; n_Control_ = 85). In order to be included in the comorbid group, the participant had to have a confirmed diagnosis of either GAD or MDD, comorbid with any other disorder(s) (see Table S1).

### 2.2 Experimental Procedures

Upon entering the lab, informed consent was obtained after which participants completed a battery of cognitive tests and questionnaires. Cognitive tests included the Rey-Auditory Verbal Learning Test (RAVLT), Trail Making Test parts A and B, Digit Span forward and backward, Controlled Oral Word Association Test, and animal fluency. The State Trait Anxiety Inventory (STAI), Penn State Worry Questionnaire (PSWQ), and Beck Depression Inventory-Second Edition (BDI-II) were administered as measures of psychiatric symptom severity. All measures collected are reported here for the sake of transparency; however only the BDI-II, STAI, and PSWQ, measures commonly used in clinical settings for psychopathology symptom quantification, were used in the data analyses of the current paper. Therefore, no further information will be reported on the other measures (see Baldwin et al., 2015 & Clawson et al., 2013 for comprehensive information). Information on the psychometric properties of the measures included is reported below in section 2.3.

Following completion of the neuropsychological tests and symptom questionnaires, participants completed a modified arrow version of the Ericksen flanker task (Eriksen & Eriksen, 1974). Participants were presented with five arrows and asked to respond to the direction of the point of the middle arrow with an index or middle finger button press. There was a total of 798 randomly presented trials with 354 trials (45%) being congruent (e.g., <<<<<) and 444 trials being incongruent (55%) (e.g., <<><<). Participants completed a practice block of 24 trials prior to the beginning of the task to ensure understanding. Stimuli were presented in white 36-point Arial font on a black background on a 17-inch computer approximately 20 inches from the participant. Flanking arrows were presented for 100 ms followed by the target arrow which was presented for an additional 600 ms. Subsequently, a fixation cross was shown for a jittered intertrial interval of 800, 1000, or 1200 ms. Responses occurring over 1600 ms after stimulus presentation were seen as an error of omission and were not included in the data analyses as the next trial was queued after 1600 ms.

### 2.3 Measures

Means and standard deviations along with Chronbach’s alpha (overall and by group) are presented in Table 1. The Beck Depression Inventory, Second Edition (BDI-II; Beck, Steer, & Brown, 1996) was used to quantify depressive symptoms. Participants were asked to rate 21 statements on a scale from 0 (I do not feel sad) to 3 (I am so sad or unhappy that I can’t stand it) after which individual item scores were summed to a total score. Possible scores range from 0 to 63. The BDI-II has been shown to have a high level of internal consistency (Chronbach’s alpha .89-.93; Beck et al., 1996; Whisman, Perez, & Ramel, 2000).

The State Trait Anxiety Inventory (STAI form Y-2) was used to quantify trait anxiety symptoms (Speilberger, Gorsuch, & Lushere, 1970). Items on the STAI include statements such as “I feel calm” or “I am worried”. Participants were asked to rank the statements on a four-point Likert type scale ranging from “not at all” (1) to “very much” (4). Because only trait anxiety is of interest to the current study, we just used trait anxiety subscale score was used for analyses. Possible scores on the STAI trait subscale range from 20-80. In previous studies, the STAI shows good internal consistency (Chronbach’s alpha>.7; Bergua et al., 2012; Speilberger et al., 1970).

The Penn State Worry Questionnaire (PSWQ; Meyer, Miller, Metzger, & Borkovec, 1990) was used to quantify anxious apprehension and worry symptoms. Participants were presented with 16 items and asked to rank their feelings on a 5-point Likert scale from “Not at all typical of me” (+1) to “Very typical of me” (+5). Items were reverse scored as needed and total score for the PSWQ was calculated through the summing of each item. Possible scores on the PSWQ range from 0-80. The PSWQ has good validity and internal consistency (Meyer et al., 1990).

### 2.4 Electroencephalogram recording and reduction

Data were collected from 128 equidistant passive Ag/AgCl electrodes on a hydrocel sensor net from Electrical Geodesics, Inc. using a NA 300 amplifier system (EGI; Eugene, OR; 20K nominal gain, bandpass = 0.01 – 100 Hz). During data collection, all data were referenced to the vertex electrode (Cz) and digitized continuously at 250 Hz with a 16-bit analog to digital converter. Per the manufacturer’s recommendation, impedances were kept at or below 50 kΩ. Offline, all data were digitally high-pass filtered at 0.05 Hz filter and digitally low-pass filtered at 30 Hz in NetStation (v 5.3.0.1). Data were then segmented from −1000 ms before response until 1000 ms after response for both correct and error trials for the time-frequency analyses, and 400 ms before response to 800 ms after correct and erroneous responses for ERN analyses.

Segmentation was extended for the time-frequency analyses from the traditional ERN segmentation in order to create a long enough epoch to extract low delta frequencies and to avoid edge artifacts common in time-frequency analyses (Cohen, 2014). For both the ERN and time-frequency measures, following segmentation eye movements and blink artifacts were corrected using independent components analysis (ICA) in the ERP PCA toolkit in Matlab (Dien, 2010). If any ICA component correlated with two blink templates (one template being provided by the ERP PCA Toolkit (Dien, 2010) and one template being derived from previous data by the authors) at a rate of 0.9 or higher, the specific component was removed from the data (Dien, 2010). Additionally, if the differential average amplitude was greater than 50 microvolts or if the fast average amplitude of a particular channel was greater than 100 microvolts, the channel was defined as bad and the nearest neighbor approach (using six electrodes) was used to interpolate the data for that electrode (Dien, 2010). Following artifact correction, data were re-referenced to an average reference in the ERP PCA toolkit in Matlab and baseline adjusted from 400 ms to 200 ms pre-response for all measures.

#### 2.4.1 Time-Frequency Data Reduction

Time-frequency power values were extracted through Matlab (R2018a) from four fronto-central electrodes (6 [FCz], 7, 106, 129 [Cz]; see Larson et al., 2014 for electrode montage). These electrodes were chosen as we were combining both time- and time-frequency domain indices of error-monitoring, and ERN amplitude is maximal over fronto-central electrodes (e.g., Clawson, South, Baldwin, & Larson, 2017). Twelve log-spaced frequencies ranging from 1.5 Hz to 14 Hz were used for a complex Morlet wavelet convolution with trial averaged EEG data. To avoid edge artifacts that are common in time-frequency analyses (Cohen, 2014), 300 ms of data were removed from the epoch, with 100 ms being removed pre-stimulus and 200 ms being removed at the very end prior to convolution. This resulted in a final epoch of 900 ms before response until 800 ms after. Due to the imbalance of correct and error trials, a random permutation of correct trials matching the number of error trials were selected for each participant (Cohen, 2014) as to not bias results towards one trial type or the other. Thus, each participant had the same number of error and correct trials for all analyses (all trial numbers per group are reported in Table 2). After, wavelet convolution was performed using complex Morlet wavelets, data were decibel baseline normalized with a condition-average from 400 ms to 200 ms prior to response. Data were then grand averaged across all groups and visually inspected to determine a time window from which to extract delta and theta power values (similar to the collapsed localizer approach advocated in Luck & Gaspelin, 2017). The time window chosen was 0 to 150 ms following response, which is consistent with previous research examining error-related neural activity (Dehaene, Posner, & Tucker, 1994; Gehring et al., 1993). Average delta power for correct and error trials was extracted from the 1-4 Hz range while average theta power was extracted from the 4-8 Hz range.

#### 2.4.2 Error-Related Negativity Data Reduction

Event-related potential values were extracted using Matlab (R2018a) and R (v. 1.1.463) from the same four fronto-central electrodes (6 [FCz], 7, 106, 129 [Cz]). After all data were baseline adjusted from 400 ms to 200 ms pre-response, mean amplitude was extracted for both error and correct trials (ERN and CRN amplitude respectively for time-domain measures) from 0 to 150 ms post-response. Mean amplitude measure was employed due to research suggesting the mean amplitude is more reliable than other ERP peak measures (Clawson et al., 2013; Luck, 2005). All means and standard deviations of dependent variables and trial numbers are reported in Table 2.

### 2.5 Data Analysis

#### 2.5.1 Questionnaire and Behavioral Data

All statistical analyses were performed in R (v 3.5.2). To determine if individuals with a diagnosis of pathology did indeed present with greater anxiety and depressive symptoms, three 4-group (GAD, MDD, Comorbid, Control) one-way ANOVAs were conducted, one for each questionnaire (BDI, PSWQ, STAI Trait) with generalized eta squared (*η*^2^) used as a measure of effect size. Post-hoc Tukey HSD were used to adjust significant group differences.

For the behavioral data, mean accuracy and median response time (RT) were calculated overall and as a function of congruency. In the flanker task, it is expected that accuracy will be lower and response time will be longer for incongruent versus congruent trials. Two 4-group by 2-congruency (congruent, incongruent) repeated measures analysis of variances (ANOVAs) were conducted with accuracy and RT as dependent variables and general eta squared (*η*^2^) used as a measure of effect size. Either paired samples *t-*tests (for within-subjects) with Cohen’s *d_z_* for effect size or follow-up one way ANOVAs with generalized eta squared for effect size were used to decompose any significant main effects or interactions

Pearson’s correlations between residualized delta/theta power and residualized ERN amplitude and all three questionnaires were conducted to characterize the relationship between all six variables.

#### 2.5.3 Continuous Linear Regressions

As a manipulation check, three paired samples *t*-tests were initially conducted on the whole sample to ensure that error trials demonstrated greater delta and theta power and more negative ERN amplitude when compared to the correct-related negativity (CRN).

In order to isolate error-related brain activity from response-related activity, the residuals between correct and error power and ERP amplitude were used as the dependent variable for the subsequent regressions. Error trials were used as the outcome variable and correct trials were used as the predictor in creation of the residualized difference scores (Meyer et al., 2017).

To test our first hypothesis that transdiagnostic measures of anxiety, worry, and depressive symptoms would predict delta power values, theta power values, and ERN amplitude, nine linear regressions were performed. In order to account for the large amount of linear regressions being performed it was decided *a priori* that only *p-*values less than 0.01 would be interpreted as significant in order to control for family wise error-rate. Age and sex were entered into linear regressions as predictors due to evidence that ERN amplitude may vary as a function of sex (Fischer et al., 2016; Hill et al., 2018; Larson et al., 2011; Moser et al., 2016) and that ERN amplitude increases as an individual ages (Tamnes, Walhovd, Torstveit, Sells, & Fjell, 2013). Each linear regression used one questionnaire as an independent variable of interest (BDI, STAI Trait, or PSWQ) to predict one dependent variable (residual delta power, residual theta power, residual ERN). Separate regressions were used as the BDI and STAI Trait scales were found to be highly correlated, and, therefore, could not be entered in the same regression. Normality of residuals was adequate.

#### 2.5.4 Diagnostic Linear Regressions

To test our second hypothesis that diagnostic group would predict greater delta residual power, theta residual power, and residual ERN amplitude, three linear regressions were performed. Group (GAD, MDD, comorbid, control), age, and sex were used to predict delta residual power, theta residual power, and residual ERN amplitude. The group variable was entered as a factored variable (i.e., dummy coded), with GAD serving as the contrast variable for each of the linear regressions.

### 2.6 Sensitivity Analysis

A sensitivity analysis performed in G*Power (v 3.1) revealed that for both the continuous and diagnostic linear regressions, the current sample is adequately powered to detect a small-to-medium *f^2^* effect. Specifically, for the continuous linear regressions, a sensitivity analysis for a linear multiple regression fixed model, R^2^ deviation from zero at an alpha level of 0.01, power of 0.80, and 3 predictors (individual questionnaire, age, sex) with a total sample size of 178 reveals sensitivity to detect an small-to-medium *f^2^* effect size of 0.09. For the secondary diagnostic linear regressions, analyses revealed that with a total sample size of 148 participants and five predictors (three diagnostic groups [with GAD set as the reference group], age, sex), we were powered to detect a *f^2^* of 0.13; both sets of linear regressions are powered to detect small to medium effects (Cohen, 1988). Thus, we are confident the results of the current study were not due to lack of statistical power.

## 3 Results

### 3.1 Questionnaire and Behavioral Data

A one-way ANOVA with BDI total score as a dependent variable revealed a difference between groups (*F*(3,142) = 35.5, *p* < .001, *η*^2^= 0.43). Individuals with psychopathology, regardless of diagnosis, had significantly higher BDI scores when compared to controls, but pathology groups did not significantly differ (*p*_GAD v Control_ < .01, *p*_MDD v Control_ < .01, *p*_Comorbid v Control_ < .01; *p*_GAD v MDD_ = .62, *p*_GAD v Comorbid_ = .06, *p*_Comorbid v MDD_ = .36). Group differences between PSWQ score were evident (*F*(3,137) = 11.1, *p* < 0.001, *η*^2^= 0.20) with individuals with psychopathology, regardless of diagnosis, having higher PSWQ scores when compared to controls and no differences amongst pathology groups (*p*_GAD & Control_ = .01, *p*_MDD & Control_ = .01, *p*_Comorbid & Control_ < .01; *p*_GAD & MDD_ = .87, *p*_GAD & Comorbid_ = .83, *p*_Comorbid & MDD_ = .26). Lastly, individuals with psychopathology had significantly higher STAI Trait scores when compared to controls (*F*(3,140) = 64.4, *p* < .001, *η*^2^= 0.58; *p*_GAD v Control_ < .01, *p* _MDD v Control_ < .01, *p*_Comorbid v Control_ < .01), but no difference between individuals with psychopathology (*p*_GAD v MDD_ = .99, *p*_GAD v Comorbid_ = .18, *p*_Comorbid v MDD_ = .08).

All behavioral (RT and accuracy) data is reported by group in Table 3. Overall accuracy for the flanker task was 91% and overall median response time was 413 ms. Paired samples *t*-tests confirmed that for the overall sample, there was lower accuracy and longer response time for incongruent trials when compared to congruent trials (*t_accuracy_*(177) = 18.6, *p* < .001, *d_z_* = 1.4; *t_response_* time(177) = 154.8, *p* < .001, *d_z_* = 11.6). For accuracy in the smaller diagnostic sample, there was a main effect of congruency (*F*(1,142) = 216.8, *p* < .001, *η*^2^ = 0.25) as expected, there was no main effect of group (*F*(3,142) = 1.63, *p* = .18, *η*^2^ = 0.03), but this was qualified by a significant group by congruency interaction (*F*(3,142) = 3.2, *p* = .03, *η*^2^ = 0.01). A paired samples *t*-test confirmed there was greater accuracy for congruent versus incongruent trials (*t*(145) = 16.3, *p* < .001, *d_z_* = 1.4). One-way ANOVAs revealed that groups did not differ on accuracy for congruent trials (*F*(3,142) = 0.43, *p* = .74, *η*^2^ = 0.01), but did on incongruent trials (*F*(3,142) = 2.70, *p* = .05, *η*^2^= 0.05). However, post-hoc tests revealed no individual group comparisons reached statistical significance (closest *p*-value = .06 between control participants and individuals with MDD). Thus, no clear differences in accuracy based on congruency and group emerged.

**Table 3.** Means and standard deviations for task accuracy <u>and response time</u>

|  | Group | Mean | Standard<br>Deviation | Range<br>(min,max) |
| --- | --- | --- | --- | --- |
| Overall flanker accuracy (%) | Overall | 91 | 6 | 57, 98 |
|  | GAD | 91 | 5 | 82, 98 |
|  | MDD | 90 | 6 | 74, 97 |
|  | Comorbid | 90 | 6 | 76, 98 |
|  | Control | 92 | 6 | 57, 98 |
| Congruent trial flanker<br>accuracy (%) | Overall | 96 | 5 | 61, 100 |
|  | GAD | 96 | 3 | 91, 100 |
|  | MDD | 95 | 5 | 75, 100 |
|  | Comorbid | 95 | 6 | 79, 99 |
|  | Control | 96 | 5 | 61, 100 |
| Incongruent trial flanker<br>accuracy (%) | Overall | 88 | 7 | 54, 99 |
|  | GAD | 87 | 7 | 74, 97 |
|  | MDD | 85 | 8 | 70, 97 |
|  | Comorbid | 86 | 7 | 73, 97 |
|  | Control | 89 | 7 | 54, 99 |
| Overall flanker RT | Overall | 413.32 | 31.68 | 317, 493.5 |
|  | GAD | 413.64 | 32.63 | 368, 477.5 |
|  | MDD | 402.54 | 33.15 | 329.5, 472 |
|  | Comorbid | 405.37 | 39.64 | 317, 455 |
|  | Control | 419.08 | 30.04 | 350, 493.5 |
| Congruent flanker RT | Overall | 371.32 | 31.93 | 278, 460 |
|  | GAD | 367.36 | 30.58 | 315, 421 |
|  | MDD | 360.14 | 34.17 | 291, 433 |
|  | Comorbid | 371.42 | 43.78 | 278, 436 |
|  | Control | 375.89 | 30.14 | 313, 460 |
| Incongruent flanker RT | Overall | 441.01 | 32.43 | 368, 525 |
|  | GAD | 446.79 | 37.67 | 401, 521 |
|  | MDD | 432.43 | 32.85 | 369, 496 |
|  | Comorbid | 431.21 | 32.78 | 368, 477 |
|  | Control | 445.68 | 31.74 | 383, 525 |

For response times in the diagnostic sample, there was a main effect of congruency (*F*(1,142) = 1660.1, *p* < .001, *η*^2^ = 0.43) and a congruency by group interaction (*F*(3,142) = 4.05, *p* = .008, *η*^2^ = 0.01), but no main effect of group (*F*(3,142) = 1.65, *p* = .18, *η*^2^ = 0.03). A paired samples *t*-test confirmed response times were longer on incongruent trials when compared to congruent trials (*t*(145) = −49.1, *p* < .001, *d_z_* = −4.1) as expected. Follow-up one-way ANOVAs revealed no differences in response times for any group for both congruent (*F*(3,142) = 1.68, *p* = .17, *η*^2^ = 0.03) and incongruent trials (*F*(3,142) = 1.93, *p* = .13, *η*^2^ = 0.04).

### 3.4 Transdiagnostic Regression Analyses

In the full sample, error trials were associated with greater delta and theta power (delta: *t*(177) = 10.1, *p* < .001, *d_z_* = 1.0, *M(SD)_error_* = 2.5(1.6), *M(SD)_correct_* =1.0(1.3) ; theta: *t*(177) = 12.5, *p* < .001, *d_z_* = 1.1, *M(SD)_error_* = 2.0(2.0), *M(SD)_correct_* = −0.1(1.6)), along with a more negative ERN amplitude when compared to correct trials (*t*(177) = −6.3, *p* < .001, *d_z_* = −0.6, *M(SD)_error_* = −0.1(2.6), *M(SD)_correct_* = 1.5(2.8)).

Scatterplots of questionnaire total score (BDI, STAI Trait, PSWQ) by each dependent variable (delta residual power, theta residual power, ERN residual amplitude) are presented in Figures 1, 2, and 3. Pearson’s correlations revealed no significant relationships between psychiatric symptoms measured by the questionnaires and error-related EEG/ERP dependent variables (see supplementary Table S3 on OSF).

**Figure 1:**
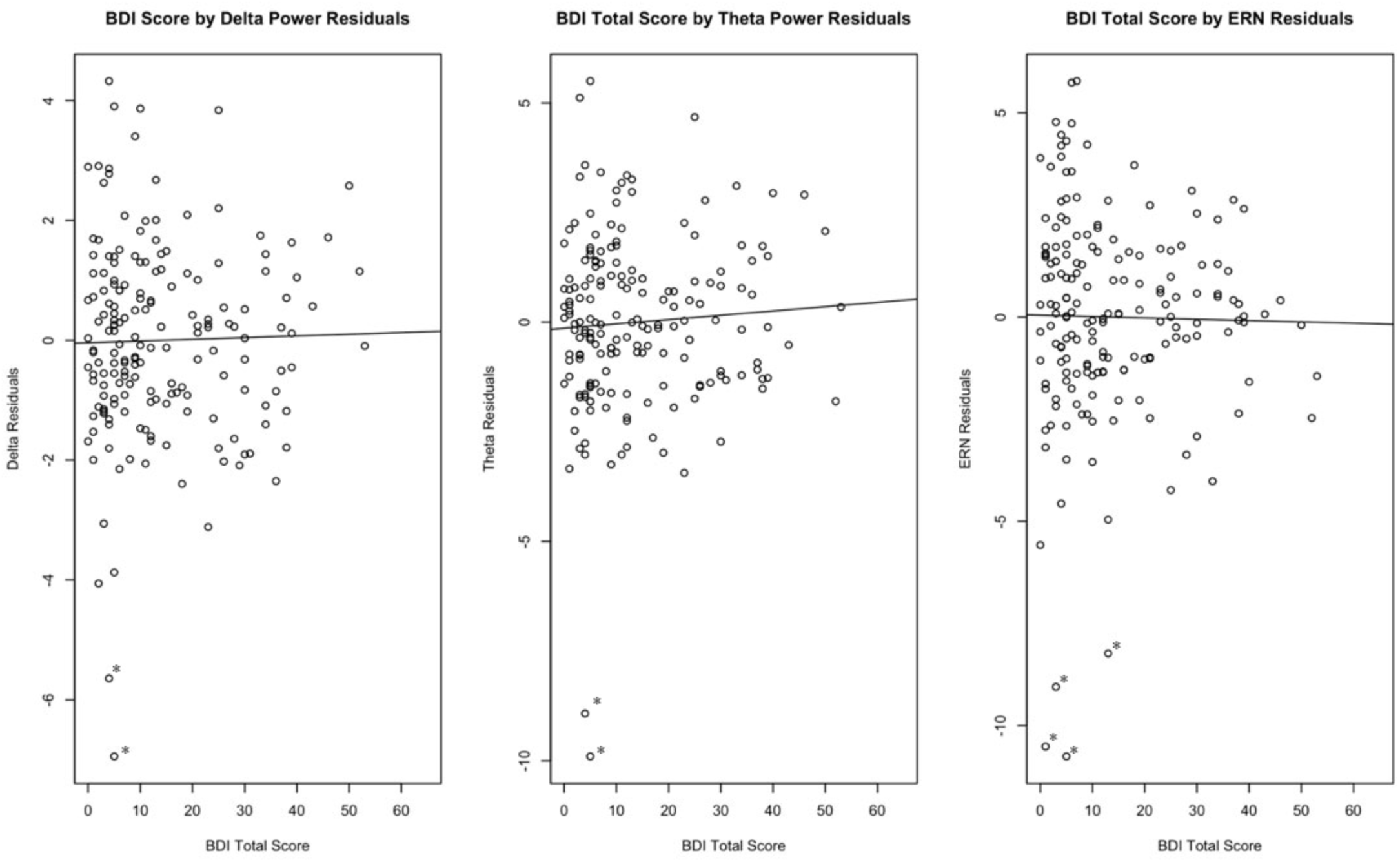
Beck Depression Inventory (BDI-II) total score by dependent variables. Possible BDI-II scores range from 0 to 60. * Outliers are marked with an asterisk. Outliers were identified as 2 times the inter-quartile range and taken out for exploratory regressions. Results did not change. All *p >* .23.

**Figure 2:**
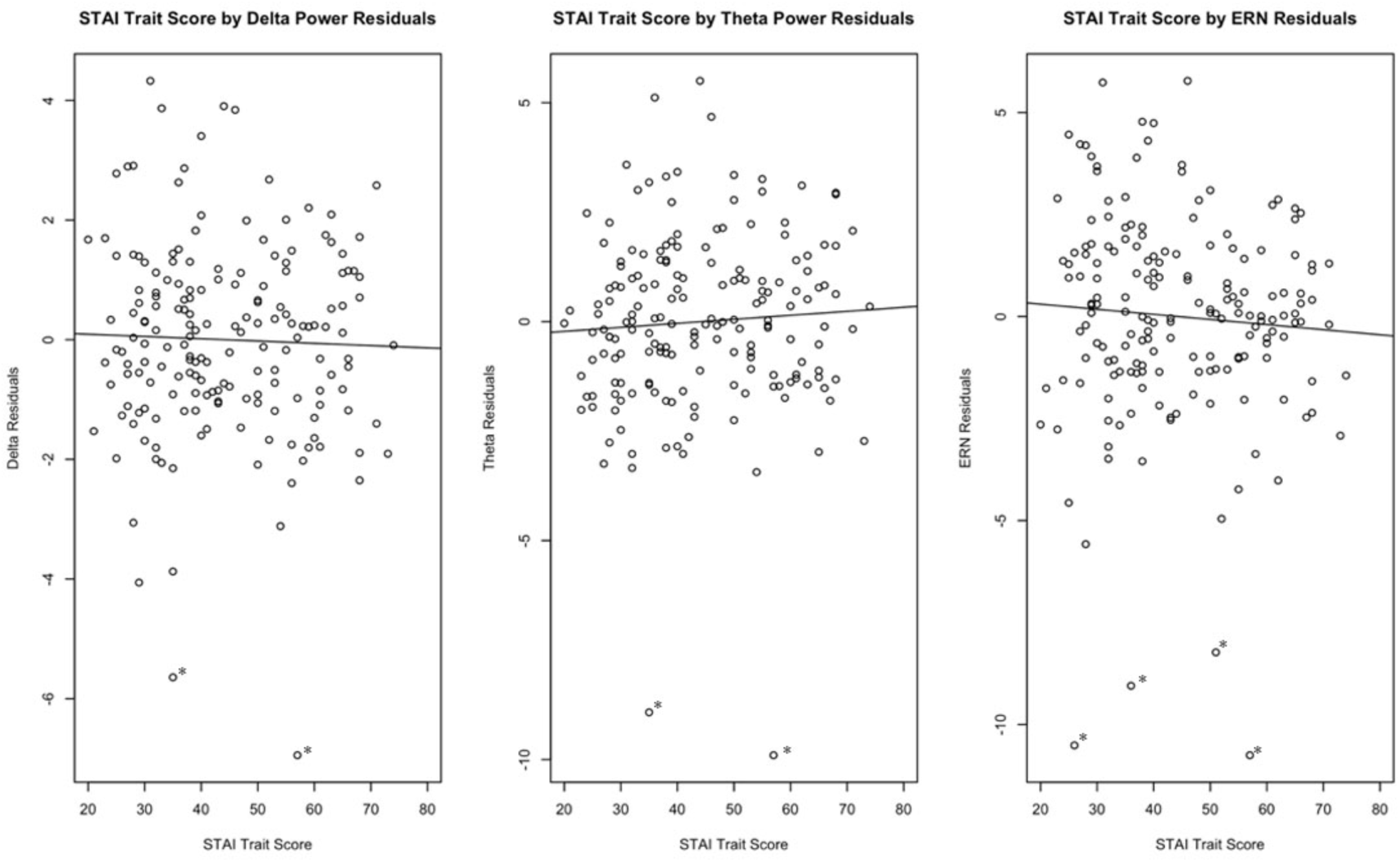
State Trait Anxiety (STAI) trait scale by dependent variables. Possible STAI scores range from 20-80.

**Figure 3:**
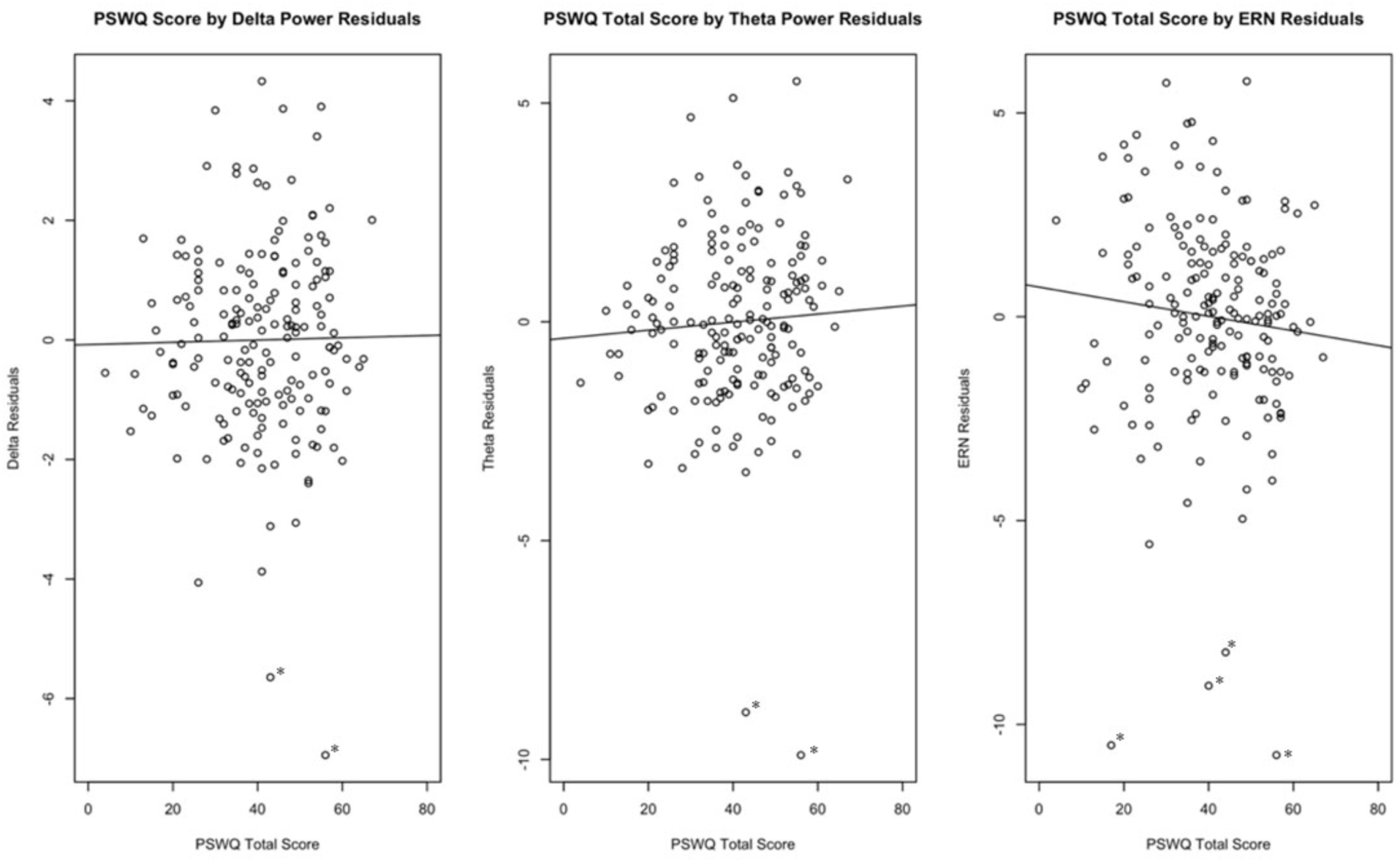
Penn State Worry Questionnaire score by dependent variables. Possible PSWQ scores range from 0 to 80.

Time-frequency plots and topographical plots for delta/theta power are presented in Figure 4 with time-frequency plots separated by group in Figure 5. Overall ERN amplitude, ERN amplitude by group, and topographical plots are presented in Figure 6. As a note, for all linear regressions, standardized beta coefficients are reported. While holding age and sex constant, BDI score, STAI trait score, and PSWQ score did not significantly predict delta power values (*β_BDI_* = 0.0, *p_BDI_* = 0.53 ; *β_STAI_* = −0.00, *p_STAI_* = 0.99; *β_PSWQ_* = 0.06, *p_PSWQ_* = 0.43; see Table 4). Transdiagnostic measures did not significantly predict theta power values (*β_BDI_* = 0.09, *p_BDI_* = 0.25 ; *β_STAI_* = 0.10, *p_STAI_* = 0.20 ; *β_PSWQ_*= 0.10, *p_PSWQ_* = 0.18; see Table 5) nor ERN residual amplitude (*β_BDI_* = −0.02, *p_BDI_* = 0.77 ; *β_STAI_* = −0.07, *p_STAI_* = 0.33 ; *β_PSWQ_* = −0.10, *p_PSWQ_* = 0.18; see Table 6).

**Figure 4:**
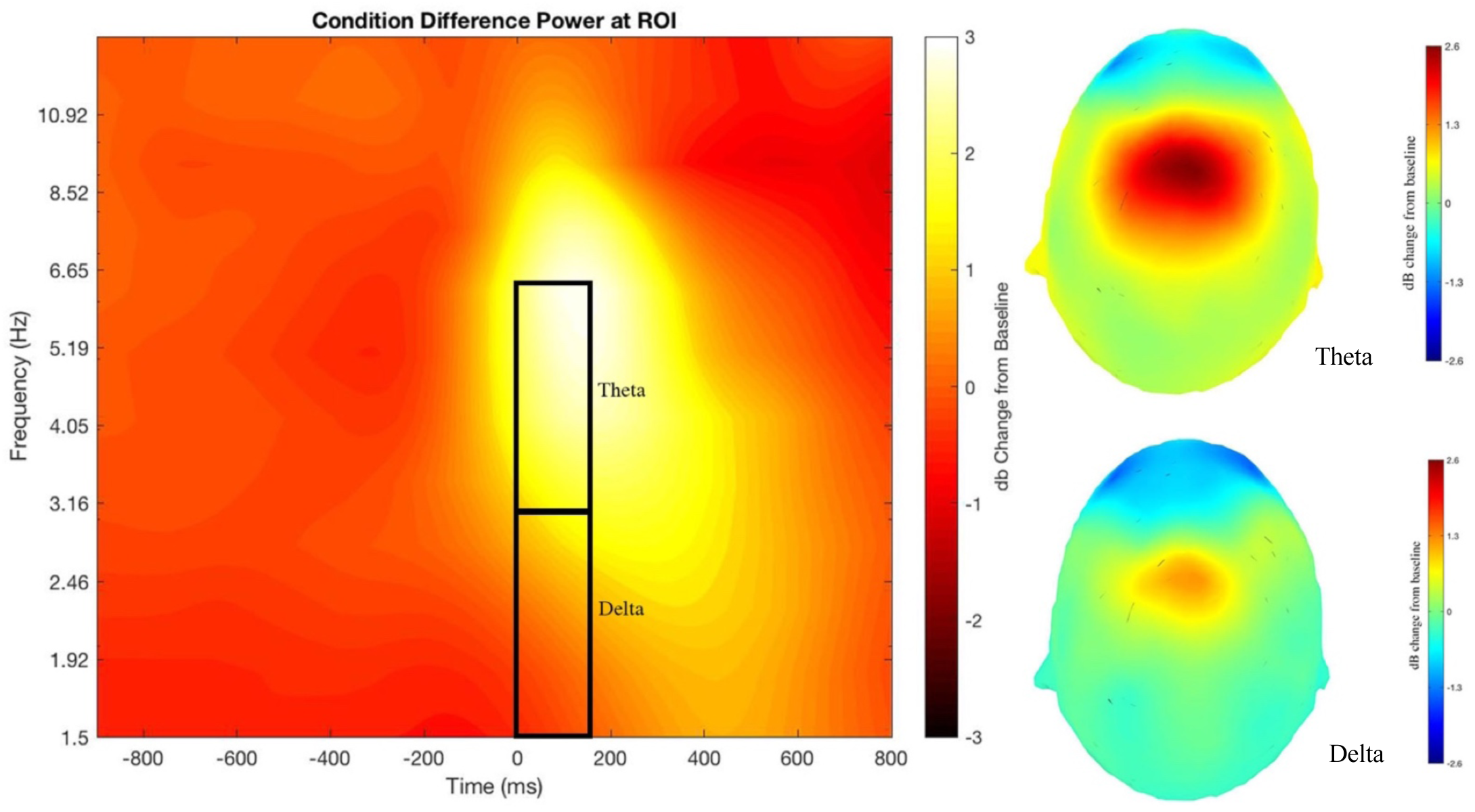
Time-frequency and topographical plots of delta and theta difference power (error minus correct).

**Figure 5:**
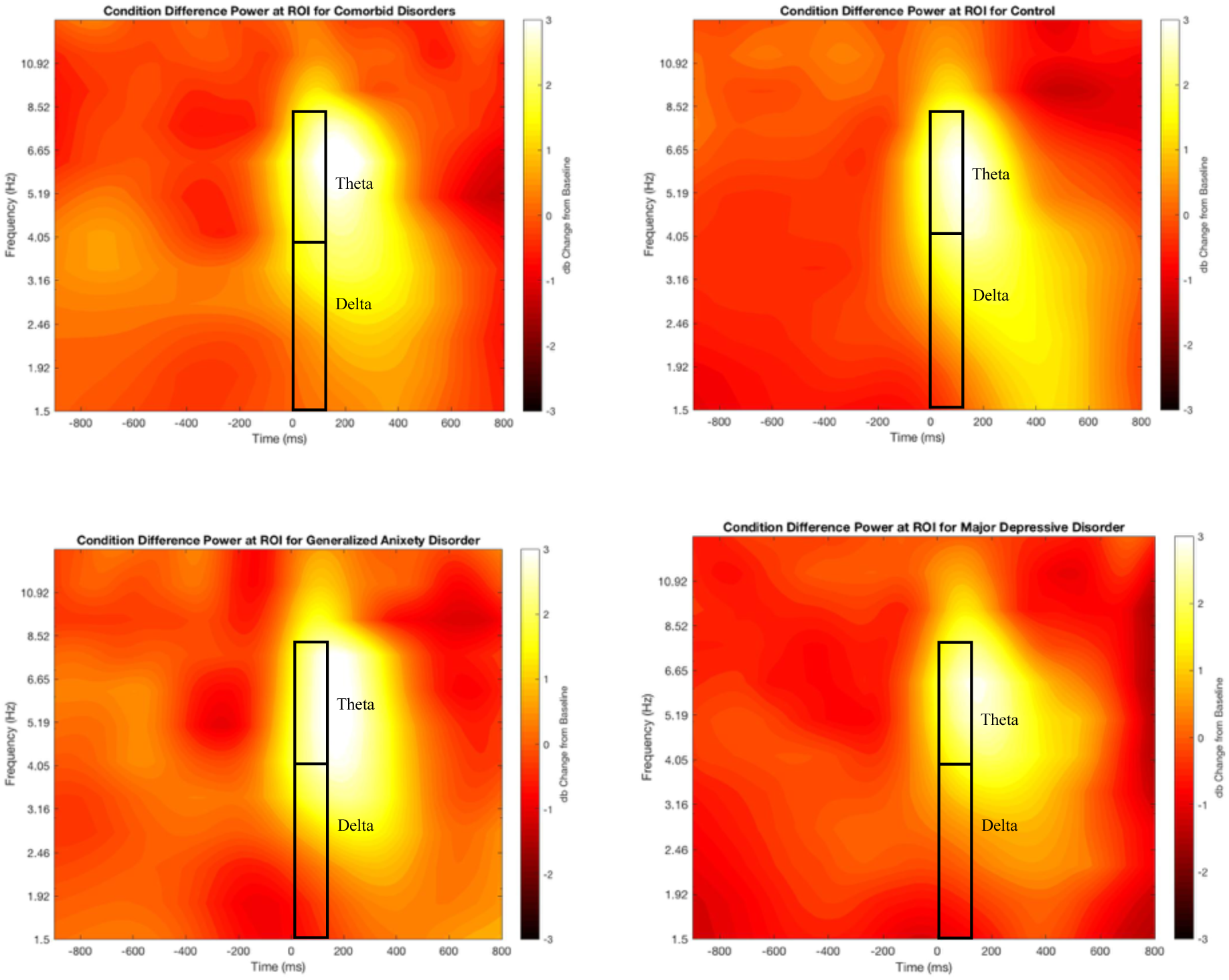
Time-frequency plots of delta and theta difference power separated by group

**Figure 6:**
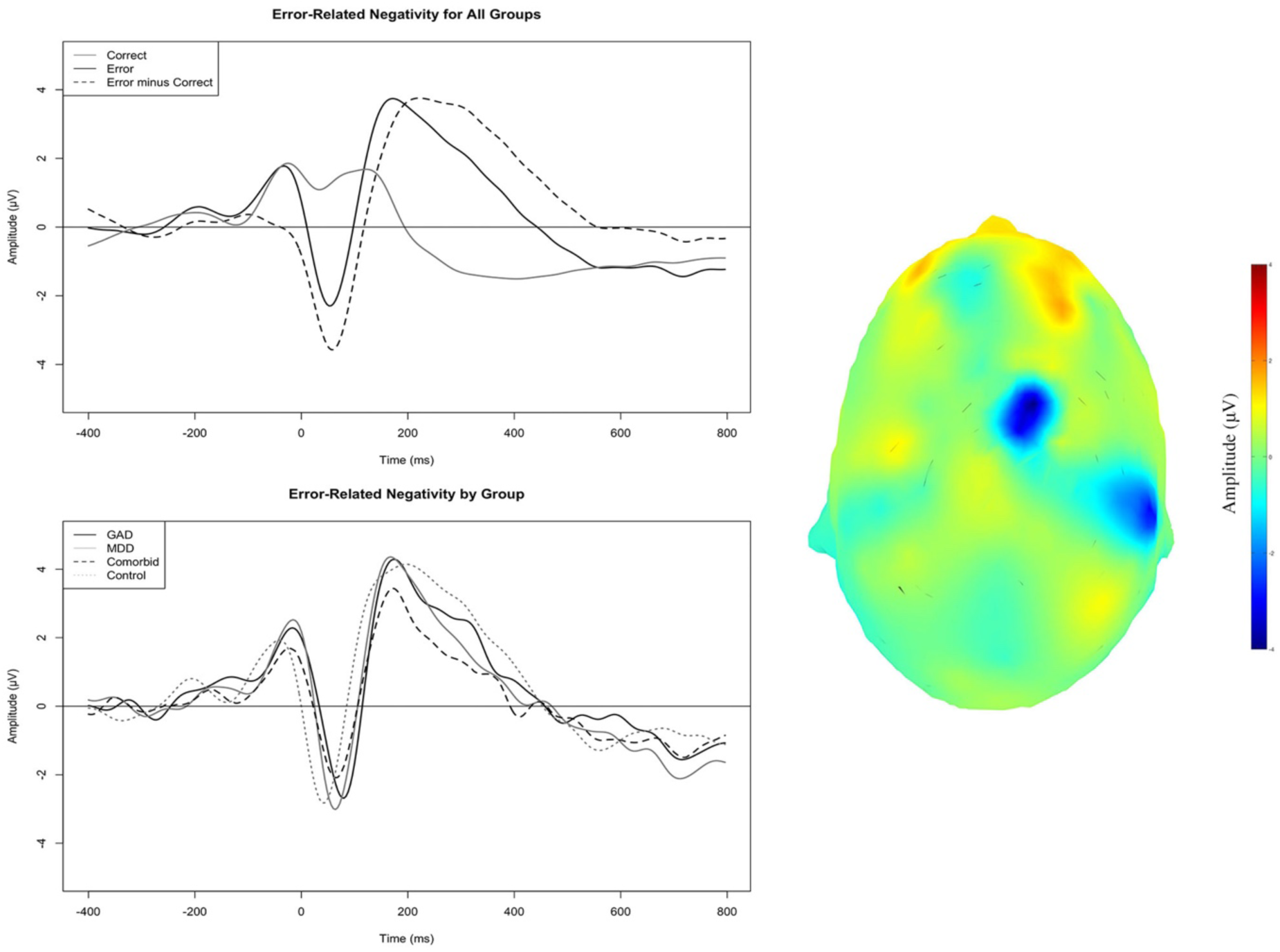
Plots of error-related negativity (ERN). Topographical plot is the difference ERN (error minus correct).

**Table 4.** Multiple linear regressions with diagnostic group predicting delta power residual values

| | $\beta$ | $t$ | $\Delta R^2$ | VIF | $F$ | $df$ | Adj. $R^2$ | Cohen's $f^2$ |
| --- | --- | --- | --- | --- | --- | --- | --- | --- |
| **BDI** |  |  |  |  | 2.9 | 3,174 | 0.03 | 0.05 |
| BDI | 0.05 | 0.63 | 0.00 | 1.02 |  |  |  |  |
| Age | -0.01 | -0.19 | 0.00 | 1.07 |  |  |  |  |
| Sex | -0.22 | -2.88** | 0.05 | 1.08 |  |  |  |  |
| **STAI Trait** |  |  |  |  | 2.76 | 3,174 | 0.03 | 0.05 |
| STAI |  |  |  |  |  |  |  |  |
| Trait | -0.00 | -0.01 | 0.00 | 1.03 |  |  |  |  |
| Age | -0.01 | -0.14 | 0.00 | 1.09 |  |  |  |  |
| Sex | -0.22 | -2.78 | 0.04 | 1.10 |  |  |  |  |
| **PSWQ** |  |  |  |  | 2.98 | 3,174 | 0.03 | 0.05 |
| PSWQ | 0.06 | 0.78 | 0.00 | 1.06 |  |  |  |  |
| Age | -0.02 | -0.25 | 0.00 | 1.09 |  |  |  |  |
| Sex | -0.23 | -2.93 | 0.05 | 1.11 |  |  |  |  |

*Note:* VIF = variance inflation factor.\* $p < 0.5$ , \*\* $p < .01$ , \*\*\* $p < .001$

**Table 5.**
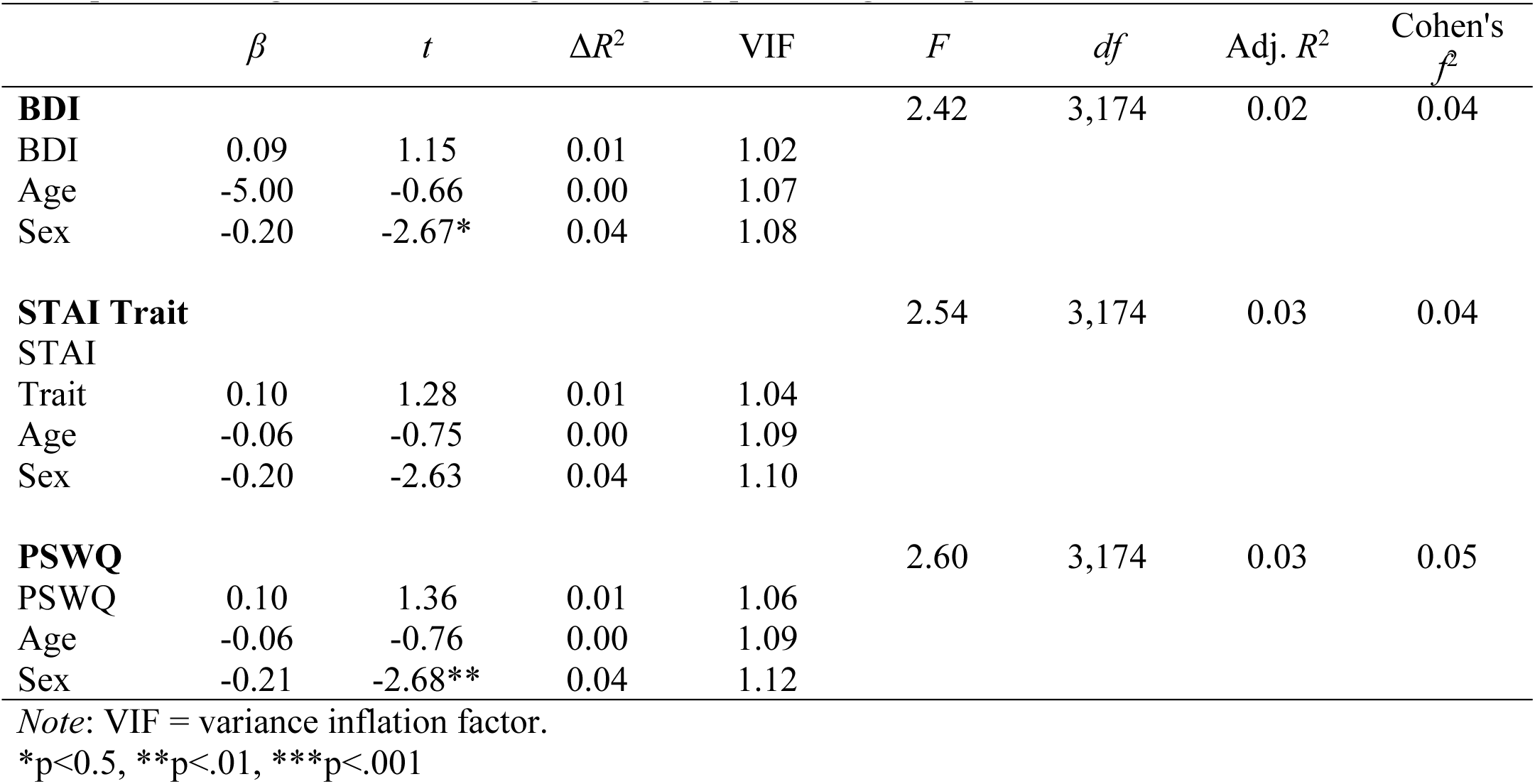
Multiple linear regressions with diagnostic group predicting theta power residual values

**Table 6.** Multiple linear regressions with diagnostic group predicting ERN residual amplitude

| | $\beta$ | $t$ | $\Delta R^2$ | VIF | $F$ | $df$ | Adj. $R^2$ | Cohen's $f^2$ |
| --- | --- | --- | --- | --- | --- | --- | --- | --- |
| **BDI** |  |  |  |  | 0.72 | 3,174 | 0.00 | 0.01 |
| BDI | -0.02 | -0.29 | 0.00 | 1.02 |  |  |  |  |
| Age | -0.06 | -0.75 | 0.00 | 1.07 |  |  |  |  |
| Sex | 0.08 | 1.02 | 0.00 | 1.08 |  |  |  |  |
| **STAI Trait** |  |  |  |  | 1.02 | 3,174 | 0.00 | 0.02 |
| STAI |  |  |  |  |  |  |  |  |
| Trait | -0.07 | -0.97 | 0.01 | 1.04 |  |  |  |  |
| Age | -0.05 | -0.63 | 0.00 | 1.09 |  |  |  |  |
| Sex | 0.09 | 1.15 | 0.01 | 1.10 |  |  |  |  |
| **PSWQ** |  |  |  |  | 1.31 | 3,174 | 0.01 | 0.02 |
| PSWQ | -0.10 | 1.35 | 0.01 | 1.06 |  |  |  |  |
| Age | -0.05 | -0.57 | 0.00 | 1.09 |  |  |  |  |
| Sex | 0.10 | 1.27 | 0.01 | 1.12 |  |  |  |  |

*Note:* VIF = variance inflation factor.\* $p < 0.5$ , \*\* $p < .01$ , \*\*\* $p < .001$

### 3.3 Diagnostic Linear Regressions

Linear regression results for the following models are reported in Table 7. While holding age and sex constant, diagnostic group did not significantly predict delta residual power (*β_MDDxGAD_* = 0.01, *p_MDDxGAD_* = 0.92; *β_ComorbidxGAD_* = 0.06, *p_ComorbidxGAD_* = 0.59; *β_ControlxGAD_* = 0.13, *p_ControlxGAD_* = 0.35). Similarly, diagnostic group did not significantly predict theta residual power (*β_MDDxGAD_* = −0.03, *p_MDDxGAD_* = 0.79; *β_ComorbidxGAD_* = 0.05, *p_ComorbidxGAD_* = 0.69; *β_ControlxGAD_* = 0.03, *p_ControlxGAD_* = 0.69). Diagnostic group did not predict ERN residual values (*β_MDDxGAD_* = 0.00, *p_MDDxGAD_* = 0.99; *β_ComorbidxGAD_* = −0.03, *p_ComorbidxGAD_* = 0.78; *β_ControlxGAD_* = 0.03, *p_ControlxGAD_* = 0.82).

**Table 7.**
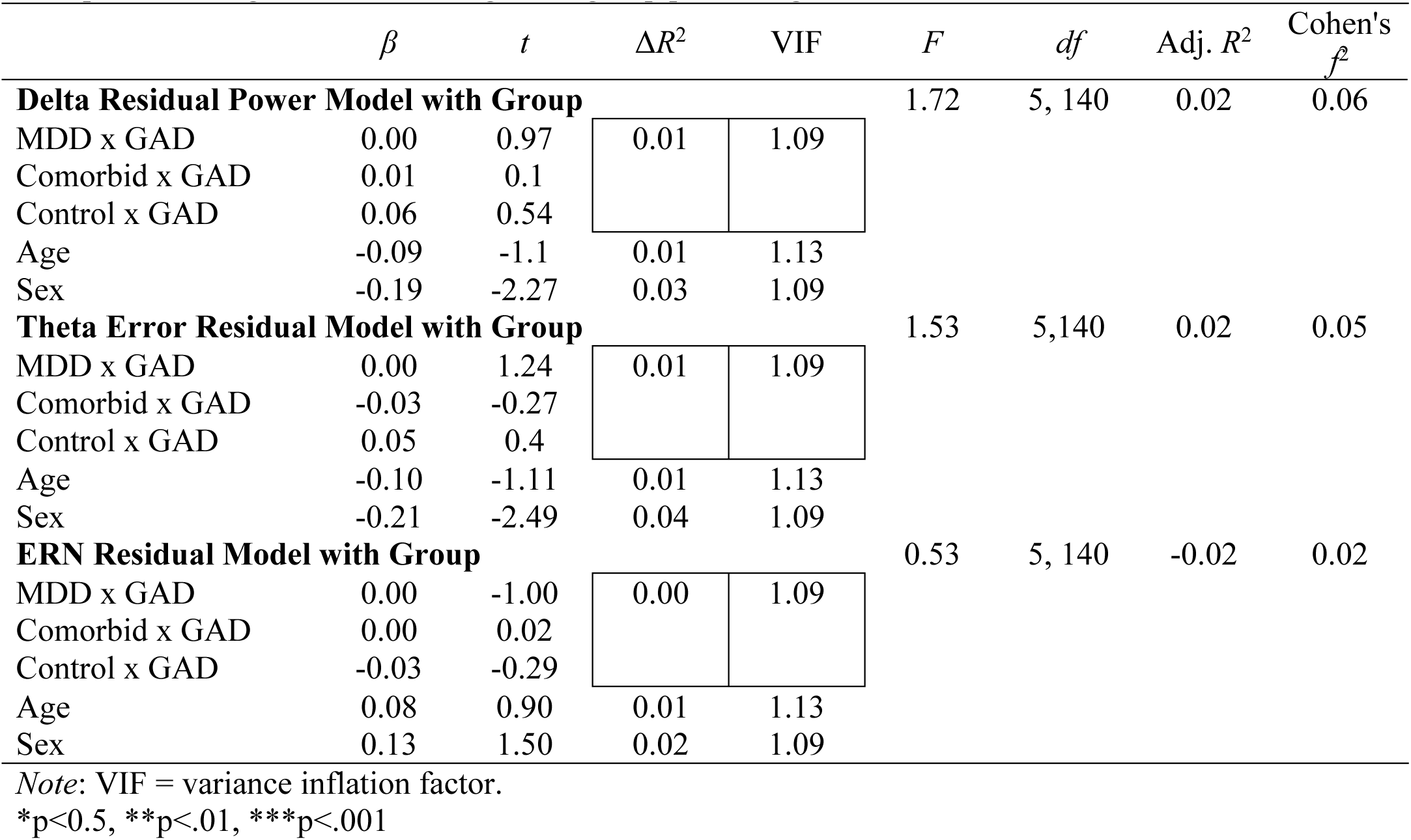
Multiple linear regressions with diagnostic group predicting residual values

Exploratory linear regressions that mirrored the regressions described above were performed with error trial only delta/theta power and ERN amplitude (i.e., not the residualized difference scores, but the error trials only). The results for these linear regressions are presented in the supplementary material on OSF (see supplementary material Tables S4-S7). All results mirrored the results presented above, with no questionnaire nor diagnostic group predicting delta/theta power and ERN amplitude. In addition, upon visual inspection of the data, there may have been potential outliers in the BDI, STAI Trait, and PSWQ scales. Therefore, to ensure that outliers were not driving the current results, outliers were defined as 2 times the inter-quartile range and taken out for exploratory regressions. The pattern of significance in the results did not change with the removal of these outliers. All *p-*values were above .23.

## 4 Discussion

The primary aim of the current study was to test the relationship between transdiagnostic measures of trait anxiety, worry, and depressive symptomology and neurophysiological measures of error-monitoring as indexed by residualized delta/theta oscillatory power and residualized error-related negativity amplitude. Our first hypothesis that higher trait anxiety and worry would predict greater residual delta/theta power and ERN amplitude was not supported, as there was a nonsignificant prediction of the residualized values from the trait anxiety and worry questionnaires. However, our hypothesis that there would be no relationship between depressive symptoms and neurophysiological indicators of error monitoring was supported, as depressive symptoms did not predict any dependent variable. A secondary aim of the current study was to test for between-group differences in error-monitoring processes in individuals with GAD, MDD, and comorbid disorders. Our second hypothesis that individuals with GAD would exhibit higher error-related delta/theta residualized power values and residualized ERN amplitude was unsupported, as group status was a nonsignificant predictor any of delta/theta power and ERN amplitude. However, our hypothesis that those with MDD would not differ from controls was supported, as there were nonsignificant differences between those diagnosed with MDD and controls.

Although the results of the current study did not support all of our original hypotheses, these results are consistent with the considerable amount of heterogeneity emerging in the extant literature. When examining the results of continuous scales predicting delta/theta power and ERN amplitude, these null results align with the results of Weinberg et al. (2014), where trait worry did not relate to the magnitude of the ERN amplitude. Further, in Weinberg et al. (2012), Mood and Anxiety Symptom Questionnaire-Anxious Arousal (MASQ-AA) subscale score did not relate to error-related brain activity, suggesting that general physiological anxiety symptoms may not be related to ERN amplitude. Although anxious arousal was not directly tested in the current study, this evidence lends credence to a general idea that anxiety symptomology and ERN amplitude may not be related without accounting for additional factors that may influence ERN amplitude, such as intolerance of uncertainty (Jackson, Nelson, & Hajcak, 2016). In addition, when looking at depressive symptoms, anhedonic depression symptoms do not relate to neural measures post-error (Schroder et al., 2013), along with general depressive symptoms (Chang et al., 2010), and distress/misery latent factors (Gorka, Burkhouse, Afshar, & Phan, 2017). Again, these results in combination with the current results suggest depressive symptoms may not be related to delta/theta power and ERN amplitude.

It is plausible there is simply not a strong relationship between anxious and depressive symptomology and neurophysiological measures of error-monitoring in a large sample of people, or that the relationship depends on extraneous variables not accounted for in the current study (i.e., hidden moderator explanation). A recent meta-analysis showed the relationship between depression and the ERN in the published literature is small (Moran et al., 2017), while another meta-analysis displayed a “small-to-medium” effect between anxiety and ERN (Moser et al., 2013), although this may be overestimated due to publication bias (Moran et al., 2017). When looking at midfrontal theta oscillations, those with higher levels of trait anxiety do display enhanced theta power when performing cognitive control tasks (Cavanagh & Shackman, 2015), but this may be specific to the individual’s reactivity to uncertainty or threat. The current results add to a heterogenous body of literature and present evidence that in a relatively large sample with a wide range of psychopathology symptoms, the relationship between transdiagnostic measures of anxiety and depression and neural indices of error-monitoring may be more nuanced than originally thought.

Another possible explanation of the current results is that error-monitoring processes may be related to more nuanced anxiety and depressive symptoms that were not captured in the measures used. We chose broad symptom measures that are commonly used in clinical settings instead of focusing on specific subscales or traits, such as anhedonia, helplessness, or rumination to name a few. It may that the relationship between neurophysiological measures of error processing and pathology are only present in very specific subdimensions. For example, individuals who experience feelings of helplessness display greater ERN amplitude when compared to those who report lower levels of helplessness (Pfabigan et al., 2013) or rumination is correlated with a more negative ERN when compared to those lower on scales of rumination (Tanovic, Hajcak, & Sanislow, 2017), suggesting that specific factors of depressive symptomology may contribute to individual differences in error-monitoring processes. Future research should continue to test which specific dimensions of depressive and anxious symptomology factors relate to error-monitoring processes in order to parse apart relationships with individual differences in error-monitoring.

When comparing the results of the group linear regressions to previous research, there is additional evidence that diagnostic group may not specifically relate to error-monitoring processes, along with methodological differences that may contribute to heterogeneity in the literature. When testing for group differences in error-monitoring processes, Kujawa et al. (2016) and Xiao et al. (2011) found no difference in ΔERN (error minus correct ERN amplitude) in individuals with GAD when compared to controls, suggesting that error-monitoring processes may not be heightened in those with GAD. However, individuals diagnosed with social anxiety disorder had a more negative ΔERN when compared to controls, suggesting that ERN amplitude may be differentially affected between anxiety disorders (Kujawa et al., 2016). Other studies have demonstrated that ΔERN was more negative in GAD, but ERN or CRN alone was not (Weinberg et al., 2012). In the current paper, residualized values between ERN and CRN values were used over ΔERN (Meyer, Lerner, Reyes, Laird, & Hajcak, 2017), and therefore methodological decisions, such as which ERP measure to use, could have affected study outcomes. When examining the literature surrounding depression and ERN amplitude, the results of a recent meta-analysis suggest that the relationship between depression and ERN amplitude is small and that the current literature is possibly contaminated with publication bias (Moser et al., 2017).

The results of the current study should be considered within the appropriate limitations. Although the sample size for the linear regressions containing all continuous variables was relatively large (n = 178), diagnostic group linear regressions had much smaller sample sizes for each subgroup (n_GAD_ = 14, n_MDD_ = 28, n_Comorbid_ = 19). Therefore, it is possible we did not have enough participants in each diagnostic category to detect small differences in neural measurements of error-monitoring that existed between groups. Further, as the task employed in the current study was originally designed primarily to extract the time domain ERN, the greatest epoch length we could extract surrounding a response was 1000 ms. Thus, the lowest frequency we could extract without violating the Nyquist theorem was 1.5 Hz (Cohen, 2014) although delta frequency extends as low as 1 Hz. This lower boundary may have impeded our ability to accurately quantify frequencies in the delta frequency range. Finally, the reliability of our symptom measures is quite good; however, reliability (in this case dependability) of ERN amplitude was below the commonly-accepted level .70. Thus, the lower reliability at .63 may have reduced the possible relationship between ERN amplitude and the symptom measures and should be considered.

In conclusion, for the current sample, trait anxiety, worry, and depressive symptoms were not related to error related delta/theta oscillatory power or ERN. Further, diagnostic group status did not predict error-related residual delta/theta power or ERN amplitude. It is possible the relationship between neural indices of error-monitoring and anxiety and depressive symptomology is more nuanced than original thought, therefore, future research should investigate various factors that could influence the relationship. It is also possible that there is not a large enough relationship between symptomology and neural indices of error-monitoring to bear out in a large sample. Future research should also investigate other individual difference traits that may influence error-monitoring to further understand what factors influence our ability to monitor errors and correct future behavior.

## Author Note

This research was supported by the Brigham Young University College of Family, Home, and Social Sciences.

There was no conflict of interest while conducting this study.

## Acknowledgements

We thank Issac Prows and Joseph Fair for their assistance in data collection.

